# A single domain intrabody as a novel tool to bias the subcellular trafficking of the follicle-stimulating hormone receptor

**DOI:** 10.1101/2025.02.13.638052

**Authors:** Pauline Raynaud, Juliette Gourdon, Vinesh Jugnarain, Lucille Berthet, Frédéric Jean-Alphonse, Océane Vaugrente, Camille Gauthier, Amandine Vallet, Priyanka Anujan, Delphine Naquin, Thomas Boulo, Christophe Gauthier, Danièle Klett, Yves Combarnous, Aylin Hanyaloglu, Eric Reiter, Gilles Bruneau, Pascale Crépieux

## Abstract

Intracellular variable fragments from heavy-chain only antibodies of camelids (intra-VHH) have been successfully used for their stabilizing properties to solve the 3D structure of active G protein-coupled receptors (GPCRs) bound to their cognate transducers. They also provide tools to link a given conformation of a GPCR to the signalling network engaged, thus allowing extensive structure/activity studies. Recently, they have been instrumental in tracking active GPCRs in various subcellular compartments. Here, we report the isolation and characterization of iPRC2, an intra-VHH recognizing the 1^st^ and 3^rd^ intracellular loops of the FSHR, but not of the luteinizing hormone/choriogonadotropin receptor close relative. Its expression in the cell decreases the cAMP production in response to hormone binding, and requires G*α*s for optimal interaction with the receptor. Importantly, iPRC2 increases the FSHR accumulation in the early endosomes, and consequently, diminishes its recycling to the cell surface. Hence, in contrast to previously described intra-VHH that disclose active GPCR intracellular location, iPRC2 provokes *per se* a location bias, through its ability to reroute the FSHR. Thus, it is an innovative tool to examine the functional consequences of GPCR accumulation in various sub-cellular compartments.

## Introduction

G protein-coupled receptors (GPCRs) constitute a large family of seven-transmembrane domain receptors involved in numerous physiological processes (1). As such, they represent the target of approximately one third of all therapeutic molecules approved by the Food and Drug Administration (FDA) (2,3). As extensively studied in the case of the β2 adrenoceptor (ADRB2), GPCRs are highly dynamic proteins that, upon ligand binding, adopt a range of intrinsically unstable conformations (4), that are stabilized by the subsequent recruitment of intracellular signal transduction proteins (4), such as heterotrimeric Gαβγ proteins (Gαs, Gαi/0, Gαq/11, and Gα12/13), as well as G protein-coupled receptor kinases (GRKs) and β-arrestins, among others, that ultimately launch a complex signalling network (5).

GPCR-targeting molecules exert a variety of pharmacological effects. They may act as agonists or antagonists, respectively activating or inhibiting GPCR signalling by interacting with it’s the receptor orthosteric binding pocket. Alternatively, they can be positive or negative allosteric modulators (PAMs or NAMs), which enhance or suppress signalling generally initiated at the orthosteric site by the endogenous ligand. When compared with the latter, agonists or allosteric modulators may favour specific signalling pathways with varying efficacy, a phenomenon known as biased signalling (6). Biased signalling does not only depend on the efficacy of receptor interactions with G proteins or β-arrestins, but also on the subcellular location where these interactions occur (7,8). Since GPCRs are known to traffic and signal from various subcellular compartments, a biased agonist might induce signalling from a cellular compartment that is not activated by the reference ligand (7). Such spatial bias can lead to both temporal and spatial modifications of the receptor-induced signalling network. For instance, in the case of the parathyroid hormone receptor (PTHR), persistent cAMP signalling from endosomes is induced only by the high-affinity agonist PTH_1-34_, but not by PTHrP_1–36_ (9). More recently, an engineered biased PTHR ligand, PTH-7d, was shown to promote persistent cAMP production at the plasma membrane, but not at endosomes, by stabilizing the receptor in a unique active conformation that couples to Gs but not to β-arrestins. In both *in vitro* and *in vivo* studies, this surrogate failed to efficiently activate vitamin D or to increase circulating calcium levels, processes that rely on endosomal signalling (10). In another study, it has been demonstrated that opioid receptors, when activated by peptide ligands, initiate signalling from both the plasma membrane and the endosomes. Non-peptide, cell-permeant ligands such as morphine also activate the opioid receptors and trigger signalling at the Golgi membrane in neuronal cells (11). This difference in signalling is likely involved in the therapeutic effects of the drugs. Another example lies with the fact that psychedelics therapeutic effects on serotonin 2A receptors (5-HT2AR) depend on their liposolubility, hence ability to activate intra-cellular 5-HT2A (12). Moreover, chemokine receptor 3 (CXCR3) endogenous ligands trigger distinct G protein and β-arrestin activation profiles at the plasma membrane versus endosomes, with endosomal β-arrestin signalling linked to inflammation in CD8+ T cells (13) via selective GRK recruitment (14). These studies highlight the potential to modulate GPCR signalling spatially, depending on the ligand involved. The ability to fine-tune these location-specific signalling pathways would offer significant therapeutic potential, enabling more precise targeting of desired physiological outcomes while minimizing unwanted side effects (7,8).

Intracellular variable domain antibody fragments from camelid heavy-chain-only antibodies (intra-VHHs) can mimic the stabilizing effects of signal transduction proteins at GPCRs. In 2011, Nb80 was the first intra-VHH shown to selectively stabilize an active conformation of ADRB2 suitable for crystallization (15), and since then, several other intra-VHHs have been successfully used as chaperones to stabilize active GPCRs for 3D structure analysis, leveraging the unique properties of intra-VHHs, particular well-suited to recognize conformational epitopes (16). Intra-VHHs specifically recognizing GPCR active conformations have also been used as biosensors to track active receptor intracellular trafficking, such as Nb80 for ADRB2 (17), or Nb33 recognizing the μ-opioid receptor (11).

Intra-VHHs also provide valuable insights into the relationship between receptor conformation and signalling activity. So far, such functional studies have been conducted on only six GPCRs out of the 800 known members of the GPCR family (18). For example, 18 intra-VHHs showed variable inhibitory effects on cAMP production and β-arrestin recruitment to ADRB2 (19). Another example is Nb6, which preferentially binds inactive conformations of the opioid receptors and reduces cAMP inhibition by Gαi at the ϰ-opioid receptor (20,21). In addition, two intra-VHH directed against the virally-encoded GPCR US28 were shown to discriminate its constitutively active versus CX3CL1-bound conformations (22). Finally, VGS-Nb2 was found to induce an active conformation of 5-HT2AR in the absence of ligand, thereby activating Gq-dependent signalling (23). In our laboratory, we isolated the iPRC1 intra-VHH that targets the follicle-stimulating hormone receptor (FSHR) and the closely related luteinizing hormone/choriogonadotropin receptor (LHCGR). This intra-VHH prevents full activation of Gαs and impairs G protein-dependent cAMP production (24) and steroidogenesis in a murine Leydig cell model (25).

The FSHR and LHCGR, together with TSHR, are class A GPCRs that constitute the glycoprotein hormone receptor subfamily. The FSHR plays a critical role in reproduction by regulating the growth and maturation of ovarian follicles and spermatogenesis. It is a key target in assisted reproductive technologies and a potential target for non-hormonal contraception. At the molecular level, FSHR primarily couples to Gαs, leading to enhanced adenylate cyclase-mediated cAMP production. The Gβγ subunits stimulate receptor phosphorylation by activating GRKs. The phosphorylated receptor then recruits β-arrestins 1 and 2 (26,27), which, in parallel (28) or sequentially (29) with Gαs and other G proteins, activate a complex signalling network (5). β-arrestins also mediate FSHR internalization (26,30). After internalization, FSHR traffics to endosomal compartments (31). While a secondary cAMP wave from endosomes is dominant in response to LHCGR activation, the proportion of cAMP generated from endosomes versus plasma membrane is less clear in the case of FSHR. The most part of FSHR is rapidly recycled back to the plasma membrane, while a small fraction is degraded through the lysosomal pathway (33). Interestingly, truncation of the last eight amino acids of the C-terminal region increases FSHR degradation, suggesting a role in endosomal sorting (33). Additionally, palmitoylation of three serine residues in the C-terminal region (at positions 644, 646, and 672) is also critical for receptor trafficking. Mutation of these residues to glycine impairs FSHR surface expression, recycling, and signalling, while increasing degradation via the proteasomal pathway (34).

While many GPCRs traffic to early endosomes (EE), LHCGR and FSHR have also been detected into distinct, smaller endocytic compartments, named very early endosomes (VEE) (31,32,35). These VEE localize close to the plasma membrane and are devoid of usual EE markers such as early endosomal autoantigen 1 (EEA1), Rab5 and phosphatidylinositol 3-phosphate (31). A subpopulation of VEE is positive for the adaptor protein APPL1, which interacts with the FSHR first intracellular loop (ICL1) (36). Protein kinase A (PKA)-dependent phosphorylation of APPL1 on serine 410 promotes LHCGR recycling (32) and limits LHCGR cAMP signalling at the VEE (32), but the impact of APPL1 phosphorylation on the FSHR recycling is less documented. A study with two small-molecule FSHR ligands, B3 and T1, have shown that from the VEE, APPL1 mediates both FSHR desensitization for Gs signalling and receptor recycling to the plasma membrane (37). The phosphorylation status of APPL1 on S410 determines the fate of the FSHR at the VEE, adding another layer of location bias within a cellular compartment (37).

In this study, we identified and characterized a new anti-GPCR intra-VHH, named iPRC2, which induces a location bias at the FSHR, providing a new tool to dissect the cellular events induced by a GPCR in selective subcellular compartments.

## Materials and methods

### References of the materials are provided in Supplementary Table 1

#### Cell culture and plasmid transfections

Human Embryonic Kidney 293 (HEK293A) cells, HEK293/ΔGαs cells (a cell line depleted for Gαs protein) (38), HEK293/ΔGαs-q-11/12 (39) and SEP-FSHR HEK293 cells (a cell line stably expressing FSHR fused to a superecliptic pHluorin) (37) were cultured in Dulbecco’s Modified Eagle Medium (DMEM) (Eurobio, Les Ulis, France) supplemented with 10% (v/v) foetal bovine serum, 100 IU/mL penicillin, 0.1 mg/mL streptomycin (Eurobio, Les Ulis, France) and kept at 37°C in a humidified 5% CO_2_ atmosphere. Murine Leydig Tumoral cells (mLTC-1) (40) were cultured in Roswell Park Memorial Institute Medium (RPMI) 1640 (Eurobio, Les Ulis, France) supplemented as DMEM, and kept in the same conditions. Cells at a density of 0.3.10^6^ cells/mL were transfected with pcDNA3.1 plasmid vector (Invitrogen, Waltham, Massachusetts, USA) in suspension, using the Metafectene Pro transfection reagent (Biontex Laboratories, München, Germany) following the manufacturer’s protocol, unless otherwise specified. Functional experiments were conducted with an excess of VHH plasmid DNA in comparison to the FSHR plasmid DNA transfected.

#### Phage library

The phage library was obtained after intramuscular immunisation of llama with a cDNA encoding the human FSHR (In-Cell-Art, Nantes, France), as previously described (41). Briefly, LeukoLOCK™ Fractionation & Stabilization Kit (Invitrogen, Waltham, Massachusetts, USA) was used to extract total RNAs from llama circulating leukocytes, which were reverse-transcribed as cDNA using SuperScript^TM^ III Reverse Transcriptase (Thermo Fisher Scientific, Waltham, Massachusetts, USA). VHHs cDNAs were amplified by nested PCR using Platinum^TM^ Taq DNA Polymerase (Thermo Fisher Scientific) (41). PCR products were inserted between the SfiI and NotI sites of the pCANTAB 6 phagemid vector (kindly provided by Pierre Martineau). The resulting recombinant phagemids were transformed in *E. coli* TG1 bacteria (Lucigen, Teddington, UK). Phages were produced upon addition of the KM13 helper phage (kindly provided by Pierre Martineau). The primary library contained 3.10^8^ independent phages.

#### Phage display

Intra-VHH selection was performed by phage display (**Figure 1**). The selection was carried out on membrane fragments of HEK293A cells expressing (or not the) human FSHR (hFSHR). To differentiate VHHs specific of the active and inactive conformations of the FSHR, selection was performed in the absence or presence of FSH, respectively. To optimize the exposure of the receptor intracellular regions to the phage library, cells were biotinylated at their surface before lysis in order to orient the membrane fragments (**Figure 1**). In the same line, biotinylated porcine FSH (pFSH) was used for stimulation of the receptor (see below). A selection has also been performed on the peptides corresponding to intracellular loops (ICLs) and carboxyterminal region of the hFSHR (**Supplemental Figure S3**).

**Figure 1:**
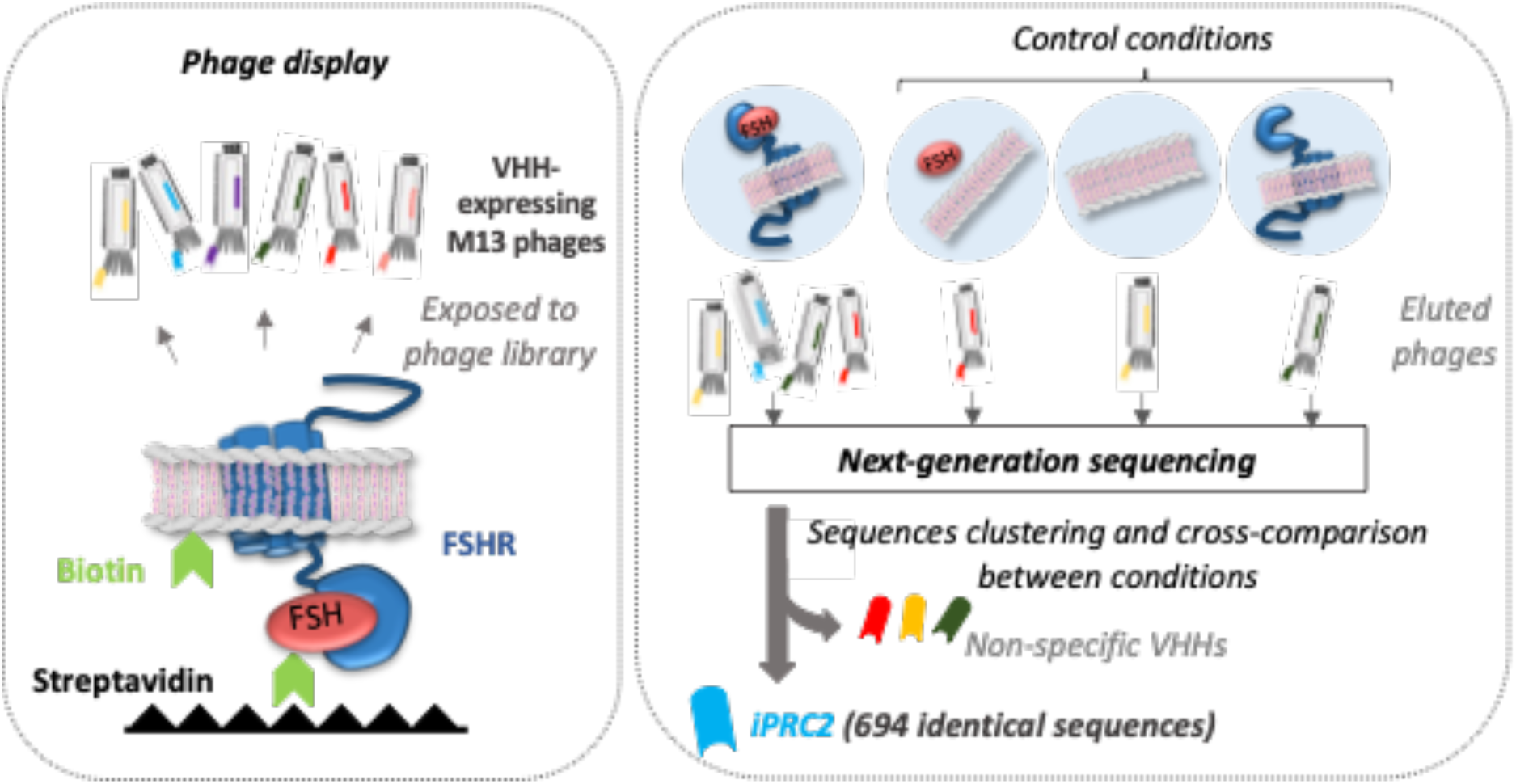
iPRC2 selection workflow. Selection of VHHs targeting the intracellular regions of the active hFSHR. Membrane fragments of hFSHR-expressing HEK293A cells stimulated with FSH were used for phage display selection. A portion of the FSH and of the cells was biotinylated prior to lysis to favour the exposure of the intracellular regions of the receptor to the phage library, using a streptavidin-biotin interaction. Control conditions included cells expressing hFSHR membrane fragments without FSH as well as mock-transfected cells membrane fragments with or without FSH. Following phage display selection, DNA encoding the VHHs from the selected phages were sequenced by NGS. For each condition, sequences from complementarity-determining region (CDR) 1 to 3 were clustered, and the sequences from each cluster were compared across all conditions. A specific cluster corresponding to iPRC2, with 694 sequences per 100,000 sequences, was identified.

*pFSH biotinylation:* porcine FSH (pFSH, CY1892-II, purified in our laboratory (42)) was biotinylated on the glycans to avoid protein alteration and to preserve receptor recognition. The chemical reaction was conducted in two steps: glycans were first functionalized through periodic oxidation. The resulting aldehydic groups reacted in a second step with the hydrazide group of a chosen biotin derivative. For the first step, sodium meta periodate (Sigma-Aldrich, St. Louis, MO, USA) at a final concentration of 4 mM was added to 140 µM pFSH in 0.1 M sodium acetate buffer pH 5.5. After a 5-minute incubation at room temperature, the mixture was immediately injected to a Fast-desalting PC 3.2/10 column (Pharmacia Biotech) and eluted with 0.1 M phosphate buffer pH=7.0 on a micro-HPLC ETTAN system (GE Healthcare). By following the absorbance at 280 nm, pFSH with functionalized glycans was collected on a Frac-950 fraction collector (Amersham), then incubated with an excess (23 mM) of biotinamidohexanoic acid hydrazide (Sigma-Aldrich, St. Louis, MO, USA) overnight at 4°C. Gel filtration using a desalting column allowed to discard the unbound biotinamidohexanoic acid hydrazide. Biotinylation of pFSH was verified by Western blotting using Streptavidin-Alexa Fluor 680 (1/5000, Thermo Fisher Scientific, Waltham, MA, USA) (**Supplemental Figure S1A**), and the ability of pFSH-biotin to induce cAMP production through hFSHR was monitored by HTRF (**Supplemental Figure S1B**).

*Preparation of cell membrane fragments:* HEK293A cells were transfected with either an empty pcDNA3.1 vector, or a plasmid encoding a N-terminally FLAG-tagged hFSHR (43) (65 ng plasmid DNA/cm^2^). Following a 2 hour-starvation in serum-free DMEM, cells were incubated with 0.1 M biotin-NHS in phosphate-buffered saline (PBS) pH 7.5 for 1 h at room temperature. To neutralize unbound biotin, cells were then incubated with 0.1 M of D-Lysine Purum for 30 minutes at room temperature, and washed 3 times with PBS pH 7.4. A portion of the cells expressing hFSHR were then stimulated with 10 nM pFSH-biotin in PBS pH 7.4 for 15 min at 37°C. For lysis, cells were scratched in 0.25 M of saccharose solution, mixed using a Polytron for 1 min and centrifuged for 10 min at 2,500 rpm at 4°C. Plasma membranes were recovered by ultracentrifugation (Beckman L8-70, rotor 70.ITI) for 30 min at 26,000 rpm. The pellet was dried and weighted before resuspension in PBS pH 7.4. Biotinylation of membrane fragments was verified by Western blot using Streptavidin-Alexa Fluor 680 (1/5000, Thermo Fisher Scientific, Waltham, MA, USA).

*Selection on a native antigen:* Three rounds of selection were performed on 100 µg of membranes (dry weight) in PBS 7.4, in Nunc Immobilizer Streptavidin C8 strips (Thermo Fisher Scientific, Waltham, MA, USA). Selection wells were either coated with membranes from cells expressing FSHR in the absence or presence of 10 nM pFSH (with 1% of pFSH-biotin), or cell membranes without FSHR, in the absence or presence of 10 nM pFSH (with 1% of pFSH-biotin) as control conditions. For the first round, a sample of 10^11^ to 10^12^ phages from the amplified phage library was used. Depletion of non-specific phages was performed on two wells coated with 100 µg of membranes from cells not expressing FSHR in the presence of 10 nM pFSH (with 1% of pFSH-biotin) for each condition and for 1 h at room temperature per well. Phages were then incubated on the selection well for 2 h at room temperature. The selection wells underwent forty washes with PBS pH 7.4, followed by three 15 min-washes in PBS pH 7.4 0.1 % Tween 20 and three 15 min-washes with PBS pH 7.4. Three more washes and a 15 min-wash were performed in 50 mM Tris pH 8.0, 1 mM CaCl_2_, before phage elution in the latter buffer containing 1 µg/mL bovine trypsin (Sigma-Aldrich, St. Louis, MO, USA), for 15 min at room temperature. TG1 bacterial cells were used for the amplification of eluted phages.

*Selection on FSHR ICLs:* Phages were selected on each of the three peptides corresponding to the intracellular loops of the human FSHR (ICL1, ICL2 and ICL3). ICL1 (TTSQYKLTVPR), ICL2 (ERWHTITHAMQLDCKVQLRH) and ICL3 (HIYLTVRNPNIVSSSSDTRIAKR) peptides were synthetized by Genecust (Boynes, France), as previously described (41).

*High throughput sequencing:* PCR products were obtained from the cDNA encoding the VHHs derived from E. coli TG1 bacteria infected with eluted phages in round 3 as templates. They were inserted into Illumina libraries, using the Next Ultra II DNA Library Prep for Illumina following manufacturer’s protocol. MiSeq Paired-End Next Generation Sequencing (NGS) was performed at the I2BC Institute (Gif-sur-Yvette, France). Sequences from the beginning of the complementarity-determining region 1 (CDR1) to the end of the CDR3 (CDRs being defined according to the IMGT reference directory) were aligned, clustered using the Hammock algorithm and compared from one selection condition to the others. Sequences common to at least 2 selection conditions were considered as non-specific, except when comparing selection against the FSHR and the ICLs.

#### Bioluminescence Resonance Energy Transfer (BRET) assays

*Interaction assay:* HEK293A cells were co-transfected with plasmids encoding the hFSHR C-terminally fused to the luciferase from the sea pansy *Renilla reniformis* (FSHR-RLuc8), AVPR2-RLuc8 or LHCGR-RLuc8 as BRET donor (0.1 μg plasmid DNA/cm^2^), and increasing quantities of plasmids encoding the intra-VHH C-terminally fused to the Venus Green Fluorescent Protein (GFP) derivative with a G4S linker as BRET acceptor (from 0 to 0.5 µg plasmid/cm^2^). To ensure an equivalent level of plasmid DNA transfected in all conditions, a pcDNA3.1 vector encoding Venus alone was co-transfected. Stimulation was performed upon addition of either 3.3 nM FSH or 100 nM AVP. The same conditions with a fixed quantity of VHH-Venus plasmid (0.2 µg plasmid/cm^2^) were used with a stimulation ranging from 0 to 33 nM of FSH to estimate interaction with inactive to active conformations of the FSHR. Further interaction experiments were conducted with an excess of VHH plasmid DNA (0.45 μg plasmid DNA/cm^2^ of either NR-Venus or iPRC2-Venus plasmid) in addition to plasmids encoding human FSHR-RLuc8, human AVPR2-RLuc8, human AVPR1A-RLuc8, human AVPR1B-RLuc8, human CXCR4-RLuc8 or human PTH1R-RLuc8 (0.1 μg plasmid DNA/cm^2^). HEK293A cells were stimulated with either 3.3 nM FSH, 100 nM AVP, 250 nM SDF1*α* (Preprotech, Neuilly-sur-Seine, France) or 10 nM PTH (Bachem, Bubendiorf, Switzerland) according to the transiently expressed receptor. For experiments in HEK293/ΔGαs-q-11/12 cells, cells were transiently co-transfected with plasmids encoding the FSHR-RLuc8 (0.1 μg plasmid DNA/cm^2^) and 0.3 µg plasmid/cm^2^ of NR-Venus or iPRC2-Venus, and stimulated with 3.3 nM FSH. For iPRC2 displacement, either 0.5 µg of an empty pcDNA3.1, or Gαs protein plasmid was also co-transfected.

*Mini-Gs protein recruitment:* for assaying mini-Gs protein (mGs, minimal engineered GTPase domain of the Ga subunit) recruitment by BRET, cells were co-transfected with plasmids encoding the human FSHR-Rluc8 (0.1 μg plasmid DNA/cm^2^), the NES-Venus-mGs (65 ng plasmid DNA/cm^2^, kindly provided by Pr. Nevin A. Lambert, Augusta University, Augusta, GA, USA) (45,46), and VHH-His (0.2 µg plasmid DNA/cm^2^). Cells were stimulated using 33 nM of FSH.

*β-arrestin 2 recruitment:* cells were co-transfected with plasmids encoding the human FSHR-Rluc8 (0.1 μg plasmid DNA/cm^2^), the Ypet-β-arrestin 2 (65 ng plasmid DNA/cm^2^, kindly provided by Dr. M.G. Scott (Cochin Institute, Paris, France) (45) and VHH-His (0.2 µg plasmid DNA/cm^2^). Cells were stimulated using a range of FSH (10^−10^ to 10^−7^ M).

*Internalisation by by-stander BRET:* cells were co-transfected with FSHR-RLuc8 (0.3 µg plasmid DNA/cm^2^) as a BRET donor, a CAAX or a LYN motif N-terminally fused to a Yellow Fluorescent Protein (Ypet) (0.06 µg plasmid DNA/cm^2^) as a BRET acceptor, and NR-His or iPRC2-His (0.2 µg plasmid DNA/cm^2^). Cells were stimulated with 33 nM FSH.

*Early endosome addressing by by-stander BRET:* cells were co-transfected with FSHR-RLuc8 (0.3 µg plasmid DNA/cm^2^) as a BRET donor, a FYVE motif N-terminally fused to a Yellow Fluorescent Protein (Ypet) (0.06 µg plasmid DNA/cm^2^) as a BRET acceptor, and NR-His or iPRC2-His (0.2 µg plasmid DNA/cm^2^). Cells were stimulated with 33 nM FSH.

*Gαs protein activation*: The ONE-GO Biosensors Kit was a gift from Mikel Garcia-Marcos (Addgene kit #1000000224) (47,48). Cells were transiently transfected with 0.1 µg plasmid DNA/cm^2^ of FLAG-FSHR, 0.1 µg plasmid DNA/cm^2^ of ONE-GO Gαs, and 0.3 µg plasmid DNA/cm^2^ of empty plasmid or corresponding intra-VHHs (pcDNA3.1-His, NR-His and iPRC2-His). Stimulation was performed with a range from 0 to 33 nM of FSH.

All BRET measurements were performed forty-eight hours after transfection, upon addition of 5 µM of coelenterazine H (Interchim, Montluçon, France) diluted in PBS for non-stimulated conditions, and with the ligand diluted in the latter buffer for stimulated conditions. Signals were recorded in a Mithras LB 943 plate reader (Berthold Technologies GmbH & Co., Wildbad, Germany).

#### Recombinant VHH production and purification

The 6xHis-tagged VHH sequences were sub-cloned in a pET22b plasmid (Novagen, Merck KGaA, Darmstadt, Germany), which was subsequently introduced in BL21DE3 competent E. coli bacterial cells (New England BioLabs, Ipswich, Massachusetts, USA). Cells were grown at 37°C and 200 rpm in 2x Yeast Extract Tryptone (2xYT) medium supplemented with 100 µg/mL ampicillin (Thermo Fisher Scientific, Waltham, Massachusetts, USA) and protein production was induced with 1 mM isopropyl β-D-1-thiogalactopyranoside (IPTG) (Sigma-Aldrich, St. Louis, MO, USA), when the culture optical density at 600 nm reached 0.5 to 0.8. Cell cultures were then centrifuged for 30 min at 4,000 *g* at 4°C. Periplasmic proteins were recovered with the osmotic shock method. The cell pellet was resuspended in 0.03 M Tris-HCl pH8.0 (MP Biomedicals, Irvine, CA, USA), 1 mM ethylenediaminetetraacetic Acid (EDTA) (Sigma-Aldrich, St. Louis, MO, USA) and 0.5 M Saccharose (Sigma-Aldrich, St. Louis, MO, USA) and the solution was stirred for 10 min at room temperature. After centrifugation at 10,000 *g* at 4°C, the cell pellet was resuspended in ice-cold 5 mM MgSO_4_ (VWR Chemicals, Radnor, Pennsylvania, USA) and stirred for 10 min on ice for periplasmic protein release. The shocked cells were centrifuged for 10 min at 10,000 *g*. Produced VHHs were purified from the culture supernatant and from the periplasmic fraction, using a Protino Ni-NTA 1 mL FPLC column for His-tag protein purification (Macherey-Nagel, Hoerdt, France), according to manufacturer recommendations. The integrity of purified VHHs was monitored by sodium dodecyl-sulfate polyacrylamide gel electrophoresis (SDS-PAGE) and thermal shift assay (TSA) (49).

#### Peptide competition by HTRF

Biotinylated peptides corresponding to FSHR ICL1, ICL2 and ICL3 (Genecust, Boynes, France) as previously defined, were used. A total of 120 µM of a biotinylated peptide were added to 30 nM purified VHHs in PBS 0.1% Tween 20 (PBS-T), and incubated overnight at 4°C and 30 rpm. Background signal was obtained with PBS-T supplemented with equivalent amount of DMSO, as in the other conditions. The sensors, MAb Anti-6His-Tb cryptate (Cisbio, Waltham, Massachusetts, USA) and Streptavidin-d2 (Cisbio, Waltham, Massachusetts, USA) were added following manufacturer protocol, and after a 1-hour incubation in the dark, fluorescence measurement was performed with a TriStar² LB 942 Multimode Microplate Reader (Berthold Technologies GmbH & Co., Wildbad, Germany).

#### cAMP measurement by Homogeneous time-resolved fluorescence (HTRF)

To quantify total cAMP in HEK293 cells, 0.5 ng Flag-tagged human FSHR (43) or 0.25 ng Flag-tagged human AVPR2 (44) plasmid DNA/cm^2^ and 80 ng of VHH-6xHis plasmid DNA/cm^2^ were used for transfection. For the condition without transfected VHH, an empty pcDNA3.1 plasmid was used. Forty-eight hours following transfection, cells were starved for 4 h in non-supplemented DMEM. Cells were detached after a 1 hour-incubation in HBSS without phenol red and Ca^2+^/Mg^2+^ at 37°C in a humidified 5% CO_2_ atmosphere. Cells were then dispatched in a 384 well-plate (Greiner Bio-one, Courtaboeuf, France), stimulated for 15 min with either 2 µM forskolin (Fsk) (MedChemExpress, South Brunswick, NJ, USA), 0.3 nM recombinant FSH (kindly donated by Merck (Darmstadt, Germany)), 0.25 nM arginine-vasopressin (AVP) (Tocris Bioscience, Noyal Châtillon sur Seiche, France) or 0.2 µM isoproterenol hydrochloride (ISO) (Sigma-Aldrich, St. Louis, MO, USA).

To quantify extracellular cAMP, mLTC-1 cells were transfected with 120 ng/cm^2^ of pCDNA.1 plasmid encoding iPRC2 or NR. After 24 hours, mLTC-1 were starved using free RPMI medium for 17 hours. Then cells were incubated with 0.1 nM LH for 3 hours, at 37°C in 5 % CO_2_. cAMP was quantified in the supernatant of mLTC-1 cells after dilution in medium.

HTRF experiments were done using the cAMP Gs dynamic kit from Cisbio (Waltham, Massachusetts, USA). After successive addition of the HTRF sensors, the plate was incubated for 1 h at room temperature. Fluorescence measurement was then performed with a TriStar² LB 942 Multimode Microplate Reader (Berthold Technologies GmbH & Co., Wildbad, Germany).

#### CRE-dependent transcription

HEK293A cells were transfected with 0.1 µg of DNA/cm^2^ of cDNA3.1 plasmid encoding FLAG-hFSHR, 0.15 µg of DNA/cm^2^ of pSOM-Luc plasmid expressing the firefly luciferase reporter gene under the control of the cAMP Responsive Element (CRE) of the somatostatin promoter region, and 0.2 µg of DNA/cm^2^ pcDNA3.1 plasmid encoding intra-VHH. After 48 h, cells were stimulated with 33 nM of FSH, for 6 h. Then, supernatants were discarded, the Bright-Glo Luciferase assay substrate (Promega, Madison, WI, USA) was added, and the emitted light was measured using a Mithras LB 943 plate reader. Values (RLU) were expressed as percentage of the maximal response obtained in the absence of VHH.

#### APPL1 Phosphorylation

HEK293 cells were co-transfected with GFP-APPL1, hFSHR-FLAG, intra-VHH-mCherry, or control mCherry plasmids, using Lipofectamine 2000 (Thermo Fisher Scientific) in accordance with the manufacturer’s protocol. After 24-48 hours, cells were harvested. For GFP-APPL1 immunoprecipitation, transfected HEK293 cells were washed three times with ice-cold PBS and collected by centrifugation at 500 × g for 5 minutes at 4°C. Cells were lysed in lysis buffer (0.5% NP-40, 10 mM Tris-HCl [pH 7.5], 150 mM NaCl, 0.5 mM EDTA, along with protease and phosphatase inhibitors) and incubated on ice for 30 minutes with gentle agitation. Lysates were centrifuged at 12,000 *g* for 15 minutes at 4°C, and the resulting supernatant was incubated with GFP-Trap agarose beads (ChromoTek) for 2 hours at 4°C with continuous rotation. Beads were then washed three times with lysis buffer and resuspended in elution buffer (120 mM Tris-HCl [pH 6.8], 20% glycerol, 4% SDS, 10% β-mercaptoethanol, 0.04% bromophenol blue). Eluted protein samples were denatured by heating at 95°C for 5 minutes before being separated on a 12% SDS-PAGE gel. Gels were either stained or subjected to Western blot analysis for further protein characterisation.

#### Total Internal Resonance-Fluorescence Microscopy (TIR-FM)

HEK293 cells stably expressing SEP-FSHR were transfected with 0.2 µg plasmid DNA/cm^2^ of either NR-mCherry or iPRC2-mCherry, using Lipofectamine 2000 (Thermo Fisher Scientific) in accordance with the manufacturer’s protocol. Twenty-four hours after transfection, cells were seeded in glass coverslips-containing dishes (Ibidi) and after 48 h, they were stimulated with 10 nM FSH. Cells were imaged using an Elyra PS.1 AxioObserver Z1 motorized inverted microscope with a scientific complementary metal-oxide-semiconductor (sCMOS) or electron-multiplying charge-coupled device (EMCCD) camera and an alpha Plan-Apochromat 100 /1.46 Oil DIC M27 Elyra objective (Zeiss), with solid-state lasers of 488 nm and 561 nm as light sources. Live cells were imaged for 1 min at 10 frames per second (fps) at 37°C in phenol-red-free DMEM supplemented with 4-(2-hydroxyethyl)-1-piperazineethanesulfonic acid (HEPES). ZEN Lite image acquisition software was utilized to collect time-lapse movies and analyzed as tiff stacks using the ImageJ plugin Time Series Analyzer. The number of recycling events counted was normalized by cell area.

#### Data analysis

Data were representative of at least 4 independent experiments and were expressed as mean values ± SD, unless otherwise specified. Data were analyzed and plotted using Graphpad Prism 9 (Graphpad Software Inc., San Diego, CA, USA). For interaction assay by BRET, nonspecific signal (when no VHH-Venus was expressed) was subtracted from the signal obtained with NR or iPRC2, and data were then normalized to NR and fitted to a specific binding curve with Hill slope (Graphpad Prism 9). For mini-Gs protein recruitment, BRET data were expressed as a percentage of the maximal FSH-induced response in the absence of VHH, and fitted to a one-phase association curve (Graphpad Prism 9). For HTRF cAMP accumulation assay, data were normalized to forskolin-induced cAMP accumulation and expressed as a percentage of ligand-induced response in the absence of intra-VHH. Statistical significance was estimated by a Mann-Whitney test to compare 2 different samples. Differences among means were considered significant at P < 0.05. Image processing was performed with Fiji (50).

#### Data availability

The nucleotide sequence of iPRC2 will deposited in Genbank upon acceptance of the manuscript

## Results

### Selection of anti-FSHR intra-VHHs

The selection was performed by phage display using an immune phage library previously described (41). To isolate VHHs directed against the FSHR intracellular side, membrane fragments from FSHR-expressing HEK293A cells were used as a native antigen. For intra-VHH selection, cell biotinylation prior to lysis was employed to favour orientation of cell membrane fragments, exposing the intracellular regions of the FSHR to the phage library, after immobilization on streptavidin-coated wells (**Figure 1**). To increase the probability of obtaining an intra-VHH recognizing the active FSHR, the selection was performed with the addition of porcine FSH (pFSH), of which 1% was biotinylated. Biotinylated pFSH was able to stimulate cAMP production in HEK293 cells expressing the FSHR, at levels comparable to those obtained with the unmodified pFSH (**Supplemental Figure S1**). As control conditions, the selection was also performed on membrane fragments from cells that do not express the FSHR, with or without pFSH (with 1% pFSH-biotin), or with the inactive FSHR (**Figure 1**). We reasoned that the greater the affinity of a phage for the antigen, the more it should be selected and amplified, and consequently, the more frequently the corresponding VHH sequences would appear. Therefore, VHHs cDNA from phages selected in each condition were sequenced by NGS, and the resulting sequences from CDR1 to CDR3 were clustered. Sequences within clusters containing at least 10 sequences were compared across all conditions. Non-specific clusters, defined as those containing a sequence that was common to at least one control condition, were excluded. This process led to the selection of iPRC2, an intra-VHH selected on the pFSH-activated FSHR. Its sequence was detected 694 times in 100, 000 sequences. An irrelevant intra-VHH not binding the FSHR (NR), previously described in (41), has been used as a negative control in all following experiments.

### The iPRC2 intra-VHH specifically binds to the FSHR, regardless of ligand binding

In this study, intra-VHHs were transiently expressed in HEK293 cells for functional characterization. Hence, we first verified that NR-mCherry and iPRC2-mCherry did not show any sign of aggregation, as assessed by confocal microscopy (**Supplemental Figure S2A**). In addition, comparable expression levels of 6xHis-tagged NR and iPRC2 was observed by flow cytometry in HEK293 cells co-expressing FLAG-tagged hFSHR (**Supplemental Figure S2B**).

The binding of iPRC2 to the FSHR in intact cells was assayed by bioluminescence resonance energy transfer (BRET) using the VHH C-terminally fused to the Venus BRET acceptor, and the receptor fused to the RLuc8 BRET donor (**Figure 2A**). As the amounts of transfected VHH-Venus plasmids increased, a dose-dependent binding of iPRC2 to FSHR was observed, which was significantly higher than the binding to the unrelated arginine-vasopressin 2 receptor (AVPR2) (**Figures 2B and 2C**). Interestingly, iPRC2 very weak binding to AVPR2 and to the closely related GPCR LHCGR was comparable (**Figures 2B and 2C**). Assessment of iPRC2 binding to various GPCRs, such as AVPR2, arginine-vasopressin 1A and 1B receptors (AVPR1A and AVPR1B), CXCR4, and PTHR, confirmed the specificity of the intra-VHH for the FSHR (**Figure 2D**). Despite our VHH selection procedure, the presence of FSH did not affect iPRC2 binding to FSHR (**Figure 2E**), suggesting that iPRC2 does not preferentially bind to the inactive or FSH-induced active conformations of the FSHR. This was further confirmed by assessing the binding of a fixed amount of NR-Venus or iPRC2-Venus transfected to FSHR in an FSH dose-response experiment (**Figure 2F**). The BRET signal provided by iPRC2 binding to the FSHR was much higher than the one obtained in the presence of the irrelevant VHH NR as expected, but no binding variation was observed upon receptor activation. To gain a more precise understanding of the iPRC2 epitope *in vitro*, HTRF experiments were achieved using purified recombinant iPRC2, in the presence of the FSHR ICLs. A positive signal was detected with ICL1 and ICL3 (**Figure 2G**). Interestingly, in an independent selection process, a specific cluster of 32 sequences per 100, 000 corresponding to iPRC2 was also selected on the ICL1 peptide (**Supplemental Figure S3**), which supports the binding of iPRC2 to ICL1 observed in this in vitro binding experiment.

**Figure 2:**
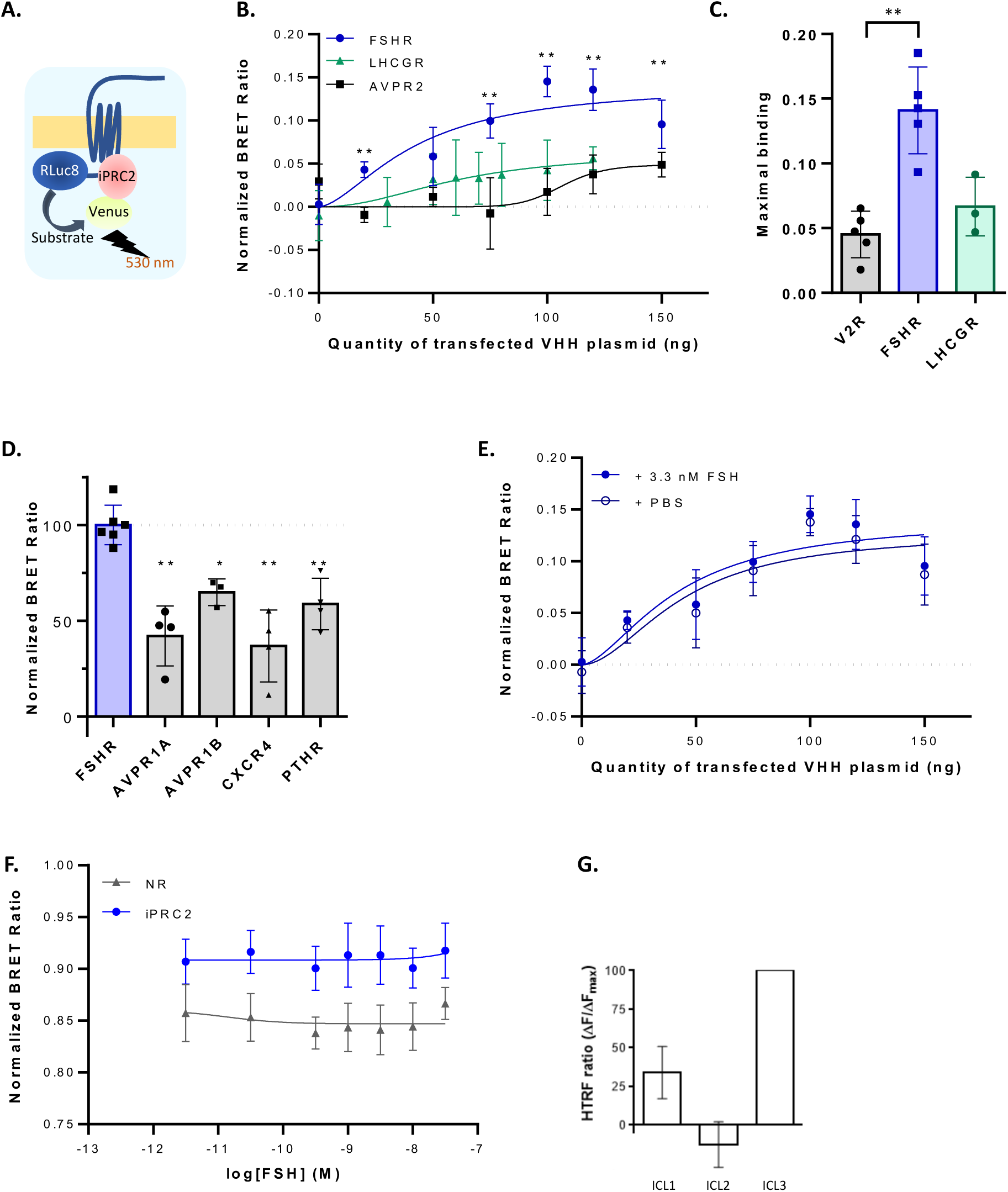
iPRC2 associates to the FSHR, whether ligand-bound or not. **(A)** Experimental design of the BRET binding assay, using FSHR fused to RLuc8 and iPRC2 fused to Venus. **(B)** BRET binding assay between FSHR-RLuc8, AVPR2-RLuc8 or LHCGR-RLuc8 and increasing amounts of transfected iPRC2-Venus plasmid. Experiments were performed in the presence of 3.3 nM FSH, 100 nM AVP or 3.3 nM LH. BRET ratios at 2 min following ligand addition were normalized to the BRET ratios obtained in the presence of the irrelevant intra-VHH (NR) fused to Venus. The curves correspond to fitted data to specific binding curves with Hill slope. The statistical significance indicated compares the FSHR to AVPR2 (N = 5). **(C)** Comparison of the maximum binding obtained for iPRC2 to the FSHR to the values obtained with AVPR2 and LHCGR (N = 5)**. (D)** BRET binding assay comparing the binding of a fixed amount of iPRC2-Venus to FSHR-RLuc8, AVPR1A-RLuc8, AVPR1B-RLuc8, CXCR4-RLuc8 or PTH1R-RLuc8. HEK293A cells were stimulated with either 3.3 nM FSH, 100 nM AVP, 250 nM SDF1*α* or 10 nM PTH, depending on the receptor. BRET ratios after 2 min of stimulation were normalized to the BRET ratios obtained in the presence of the NR-Venus intra-VHH. Statistical significance indicated compares each receptor to the FSHR (N = 4). **(E)** Same experiment as in (B) for FSHR only, comparing the curves obtained in the absence or presence of 3.3 nM FSH (N=5). **(F)** Dose-response curves of FSH to estimate the binding to the FSHR-RLuc8 with a fixed amount of transfected iPRC2-Venus plasmid or control intra-VHH (N=3). **(G)** HTRF binding assay with purified recombinant iPRC2-His to ICL1, ICL2 and ICL3 FSHR peptides (N=3). Each experiment was performed in triplicate, and data are represented as mean ± SD. Statistical significance was assessed by unpaired Mann-Whitney test, **P < 0.01, *P < 0.05.

### Functional consequences of iPRC2 binding to FSHR

Upon activation, the FSHR primarily couples to G*α*s, leading to cAMP intracellular production. Cellular expression of iPRC2 attenuated FSH-induced cAMP production by 20%, as shown by HTRF (**Figure 3A**). In contrast, iPRC2 expression had no effect on the cAMP production following activation of another Gs-coupled receptor, the β2-adrenoceptor ADRB2, endogenously expressed in HEK293 cells (**Figure 3B, Supplemental Figure S4A**). Similarly, iPRC2 had no effect on cAMP production in response to activation of LHCGR, endogenously expressed in a mouse Leydig cell line (**Figure 3C**), as expected based on our binding data. In line with these findings, iPRC2 expression also caused a significant reduction in cAMP-responsive element (CRE)-dependent luciferase activity in response to FSH, when compared to NR (**Figure 3D**).

**Figure 3:**
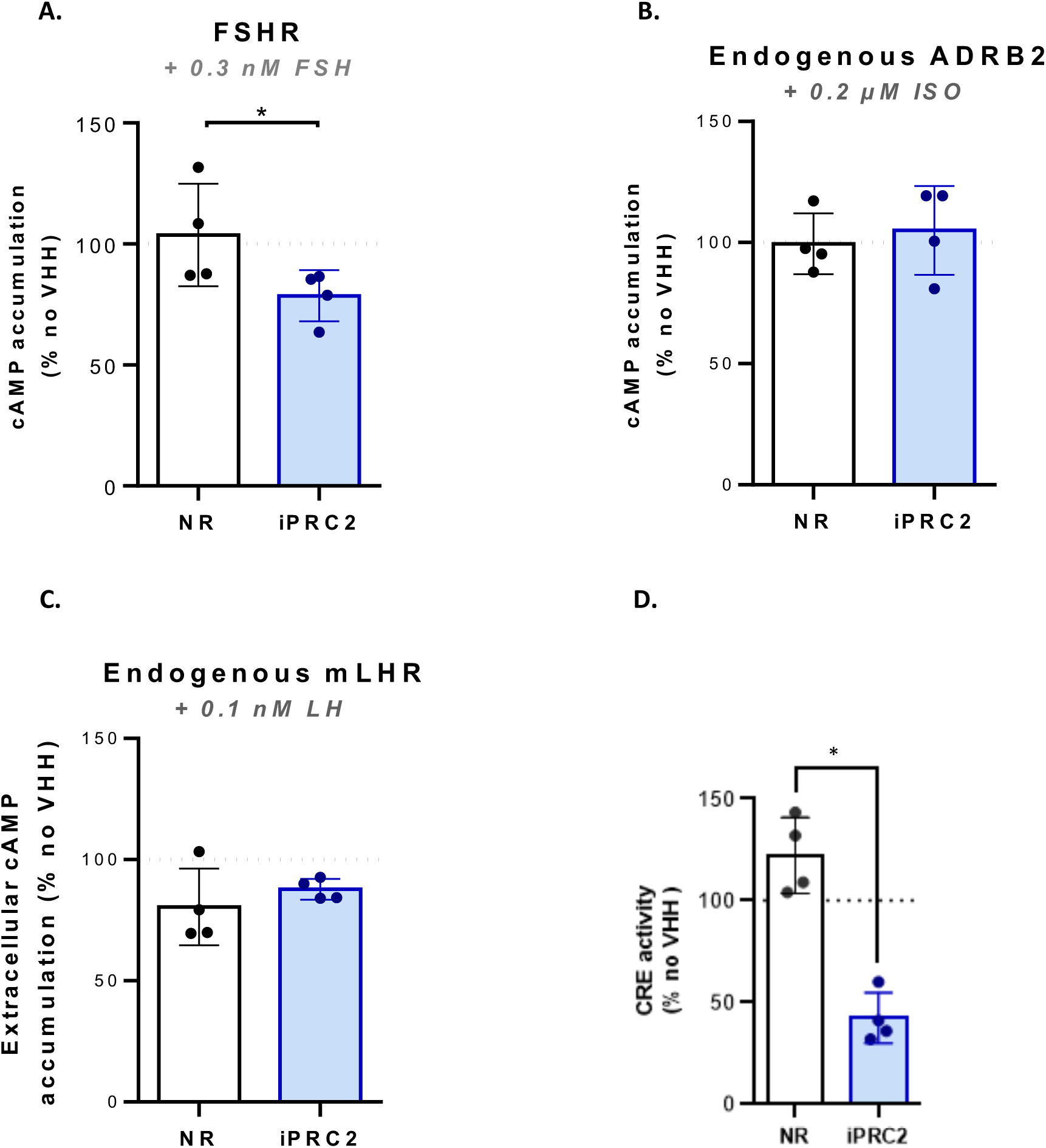
iPRC2 expression impairs FSH-induced cAMP/CRE response. **(A)** cAMP accumulation assayed by HTRF in HEK293A cells transiently expressing the FSHR. **(B)** The same experiment was performed in HEK293A cells endogenously expressing ADRB2, or **(C)** in mLTC cells endogenously expressing the mouse LHR (mLHR). Cells transiently expressed either NR or iPRC2, or were transfected with an empty plasmid (‘no VHH’ reference), and were stimulated with 0.3 nM FSH, or 0.2 µM ISO or 0.1 nM LH for 15 min. **(D)** CRE-dependent luciferase activity of HEK293A cells expressing FSHR and intra-VHH iPRC2 or NR, or transfected with an empty plasmid, and stimulated with 33 nM FSH. Data represent the mean ± SD (N=4). Statistical significance was assessed by unpaired Mann-Whitney test, * *p* < 0.05.

One possibility to explain these results is that the coupling of G*α*s to FSHR might be less efficient in the presence of iPRC2. To address this question, the recruitment of a truncated version of G*α*s (mini-Gs) to the FSHR was assessed (**Figure 4A**) in the presence of iPRC2. In parental HEK293 cells, iPRC2 did not alter mini-Gs recruitment (**Supplemental Figure S4B**). However, in cells depleted of G*α*s, a slight decrease was observed (**Figures 4B and 4C**), suggesting a possible competition. The absence of clear visible effect may be due to compensation by other G proteins that remain expressed in these cells. To further enhance the contrast, mini-Gs recruitment was assessed in cells depleted of all G*α* proteins (except G*α*i). Surprisingly, iPRC2 did not affect at all mini-Gs recruitment in this context (**Supplemental Figure S4C**). These results suggest that iPRC2 does not compete with G*α*s for interaction with FSHR. To further investigate this point, we tested whether an excess of ectopically expressed G*α*s could displace iPRC2-Venus from FSHR-RLuc8 in HEK293 cells depleted of Gs-q-11/12 (**Figures 4D**). The ectopic G*α*s protein did not displace iPCR2, but quite unexpectedly, the presence of G*α*s at the receptor appeared to enhance iPRC2 binding, an effect that was further amplified after 2, 5, and 10 minutes of FSH stimulation. This assumption was further supported by the observation that iPRC2 bound much better to the FSHR in parental HEK293 cells than in HEK293 cells depleted of Gs-q-11/12 (**Figures 4E**). Since iPRC2 did not appear to compete with G proteins for binding to the FSHR, we next investigated whether its presence affected G*α*s activation, which could potentially explain the reduced cAMP production in response to FSH. G*α*s activation was assessed using the ONE-GO biosensor kit. In this assay, a plasma membrane-anchored G*α*s-GTP detector module, containing a NLuc, generates a BRET signal in the presence of a GTP-bound G*α*s-YFP (**Figure 4F**). In this context, G*α*s GTP loading at the plasma membrane was not altered by iPRC2 (**Figure 4G**), indicating that G*α*s activity at the plasma membrane is not altered, hence not responsible for the decreased cAMP production observed in the presence of iPRC2.

**Figure 4.**
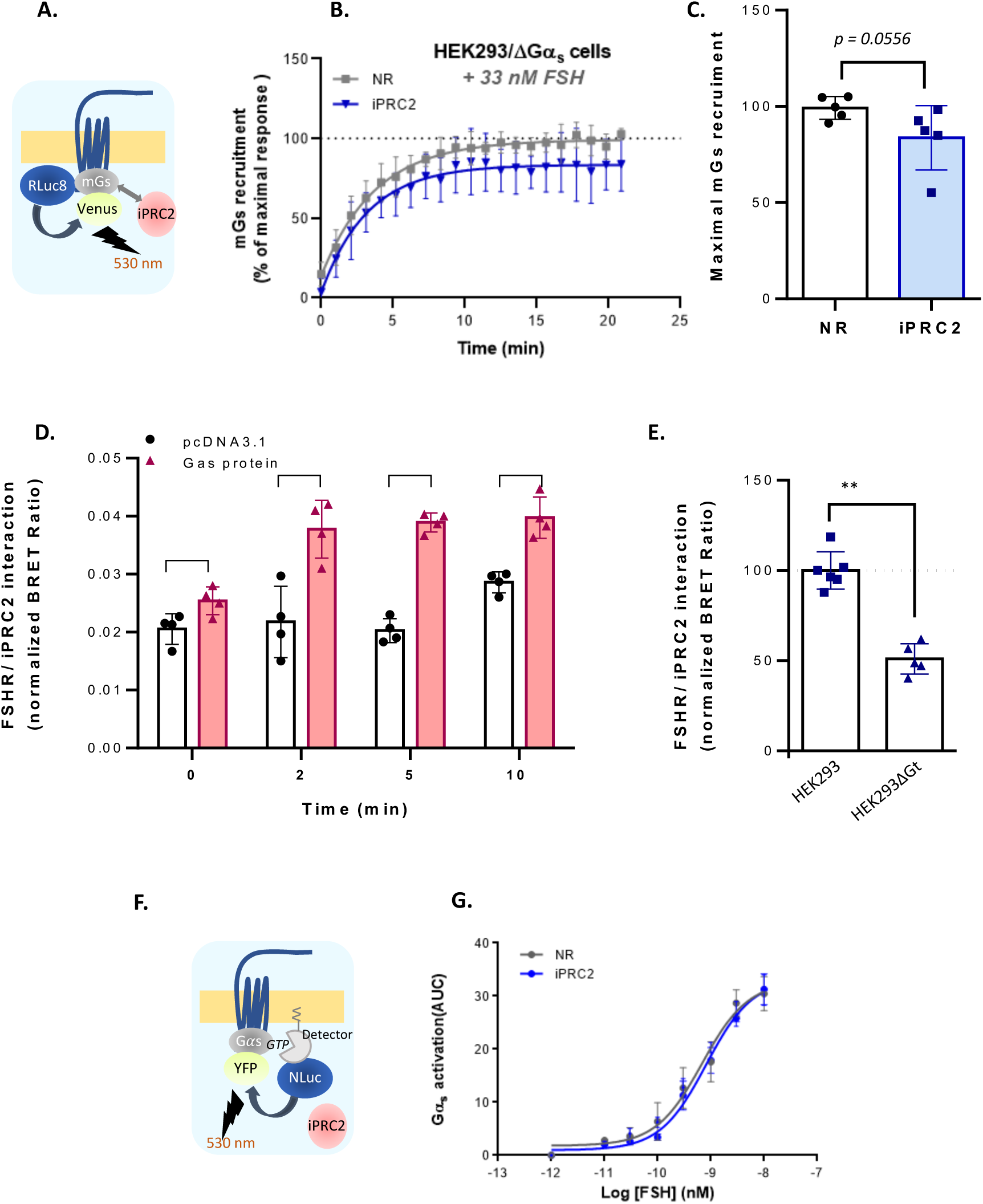
Investigating iPRC2 and G*α*s compatibility for FSHR Binding. **(A)** Experimental design of the BRET assay to monitor mini-Gs recruitment to the FSHR. **(B)** BRET assay measuring the recruitment of NES-Venus-mGs to FSHR-RLuc8, in the presence of either NR or iPRC2, in HEK293A cells depleted of G*α*s and stimulated with 33 nM FSH. The BRET ratios were expressed as percentage of the maximal FSH-response in the absence of VHH. Curves correspond to fitted data to one-phase association curves (N = 5). **(C)** The maximal recruitment of NES-Venus-mG*α*s to FSHR-RLuc8 in cells depleted of G*α*s and stimulated with 33 nM FSH was compared in presence of either NR or iPRC2 (N= 5). **(D)** Interaction assay in HEK293 cells depleted of Gs-q-11/12/ transiently transfected with FSHR-RLuc8. NR-Venus or iPRC2-Venus was transiently expressed in absence (pcDNA3.1) or in presence of overexpressed Gαs protein. BRET ratios were normalized to the BRET ratios obtained in the presence of NR-Venus (N = 4). **(E)** BRET binding assay comparing FSHR-RLuc8 and iPRC2-Venus binding, either in parental HEK293A cells or in HEK293 depleted of Gs-q/-1/12. BRET ratios were normalized to the BRET ratios obtained in the presence of NR-Venus (N=4). **(F)** Experimental design of ONE-GO biosensor kit, measuring BRET between a G*α*s-GTP detector fused to NLuc, and G*α*s-GTP fused to YFP protein. **(G)** G*α*s activation in HEK293 cells transiently expressing ONE-GO plasmids as well as FSHR, and either iPRC2 or NR was assessed with increasing concentrations of FSH (10^−12^ to 10^−8^ M). The area under the concentration/activity curves generated were plotted and fitted using non-linear regression (N=3). Each experiment was performed in triplicate, and data represent mean ± SD. Statistical significance was assessed by unpaired Mann-Whitney test.

### Consequences of iPRC2 expression on the FSHR subcellular location

The impact of iPRC2 expression on FSHR intracellular trafficking was assessed by by-stander BRET. First, FSHR internalization was measured using a Ypet fused to a CAAX motif, which interacts with the plasma membrane lipids (**Figure 5A**). No effect on FSHR internalization was observed with either NR or iPRC2 (**Figure 5B**). This result was confirmed using a different sensor, i.e., a Ypet fused to the lipid-binding domain of the Lyn protein (**Supplemental Figure S5A**). In agreement with unaltered FSHR internalization in the presence of iPRC2, β-arrestin 2 recruitment to the FSHR upon FSH stimulation was comparable when each intra-VHH was expressed (**Figures 5C and 5D**).

**Figure 5:**
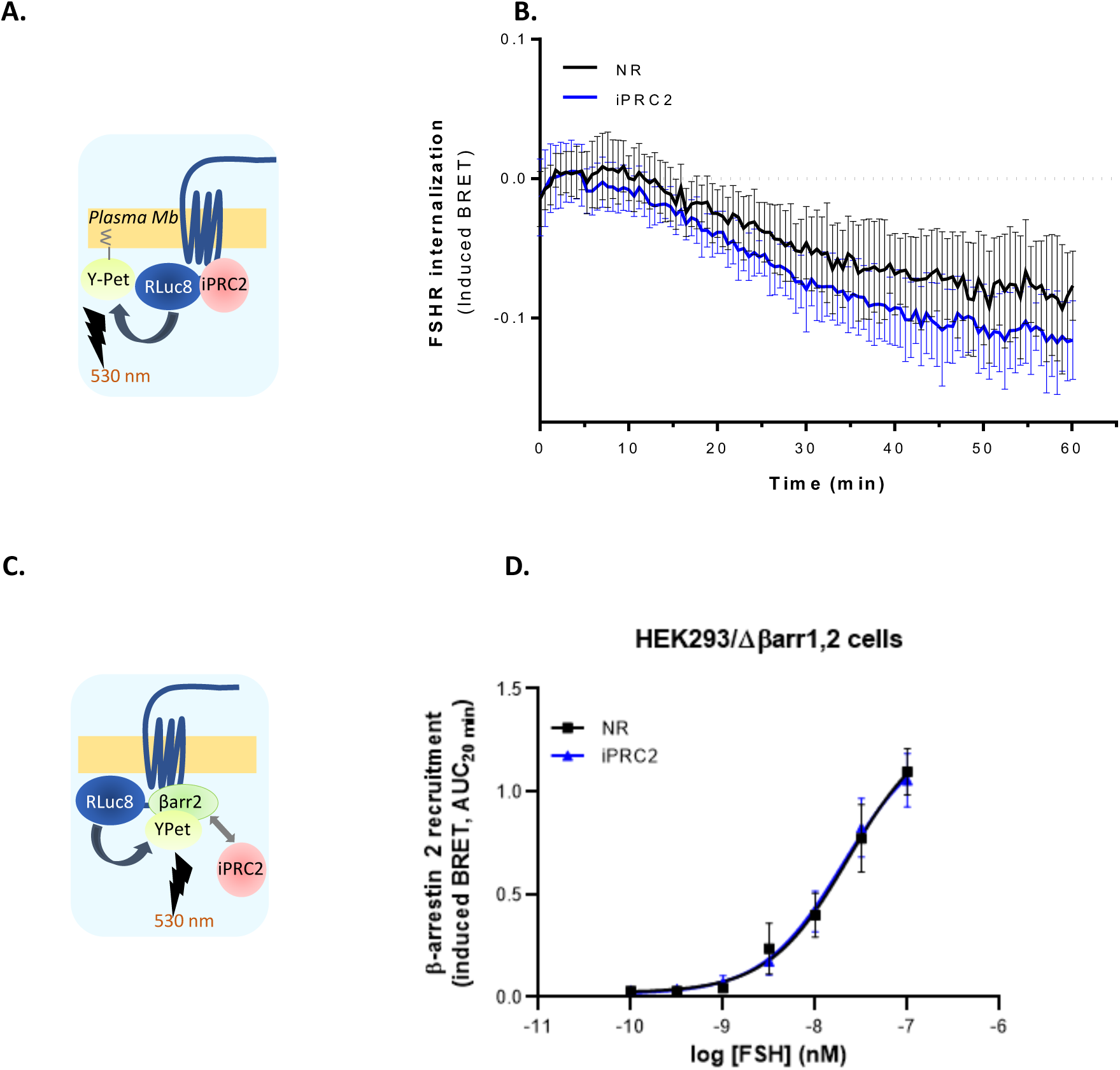
iPRC2 does not alter FSHR internalization. **(A)** Experimental design of the by-stander BRET internalization assay using FSHR-RLuc8 and a Ypet-fused CAAX motif. **(B)** Internalization assay in HEK293A cells transiently expressing the FSHR-RLuc8, CAAX-Ypet and either NR or iPRC2, after stimulation with 30 nM FSH. **(C)** Experimental design of the β-arrestin 2 recruitment BRET assay, with FSHR-RLuc8 and β-arrestin 2-YPet. **(D)** β-arrestin 2-YPet recruitment to FSHR-RLuc8 was assessed with increasing concentrations of FSH (10^−10^ to 10^−7^ M). The area under the concentration/activity curves generated were plotted and fitted using non-linear regression. All data represent mean ± SD (N=3).

However, when assayed by by-stander BRET using a Ypet fused to a FYVE motif, which interacts with the PI3P present in the EE membrane (**Figure 6A**), iPRC2 expression dramatically enhanced the accumulation of the FSHR in the early endosome (EE) (**Figure 6B**). Considering that the total number of FSHR remained constant throughout the stimulation time course, the receptor was expected to be recycled less efficiently. This was indeed observed by total internal reflection fluorescence microscopy (TIR-FM), the number of recycling events per minute being significantly reduced in the presence of iPRC2 (**Figures 6C and 6D**). Thus, iPRC2 induces *per se* a location bias of the FSHR trafficking within the cell.

**Figure 6:**
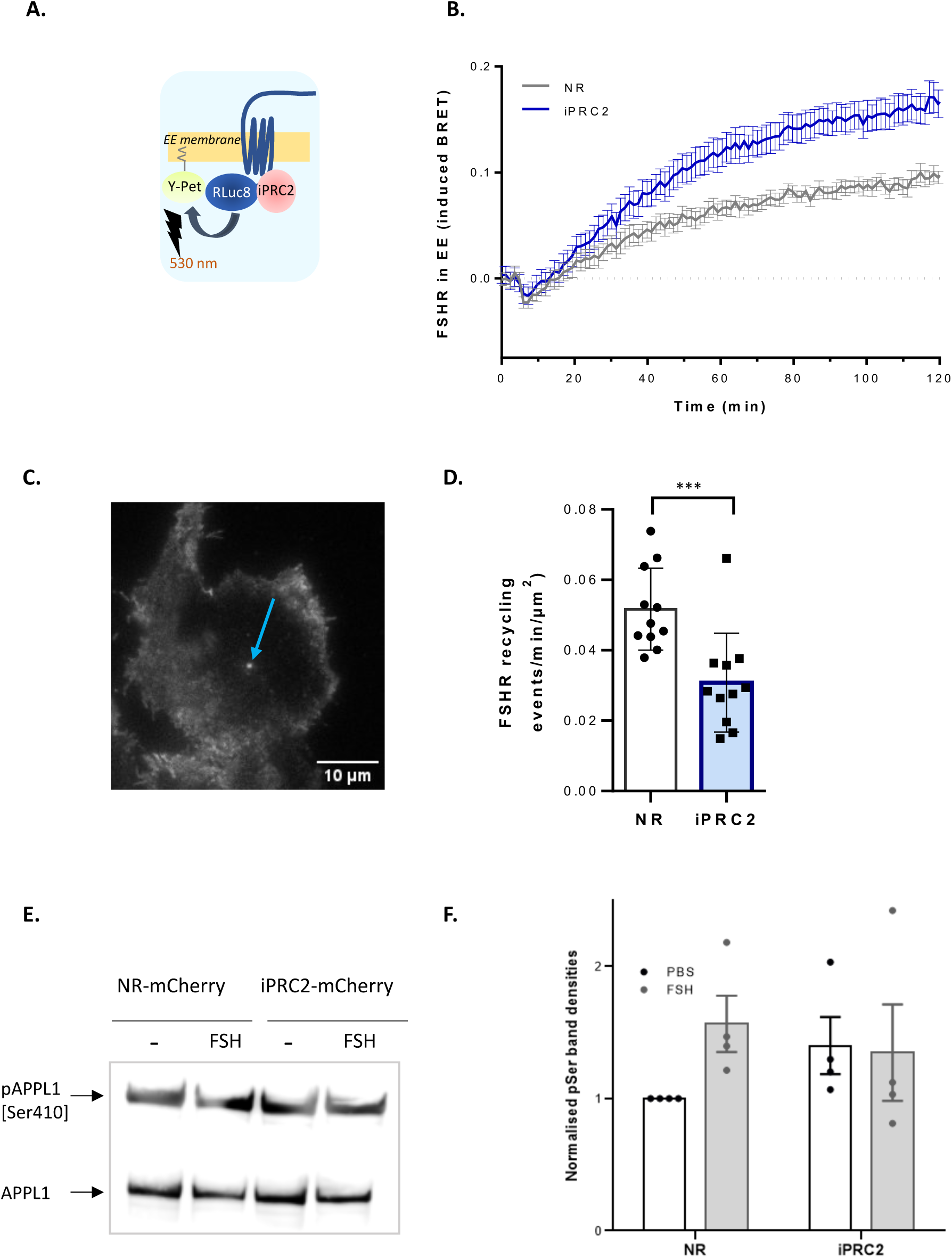
iPRC2 biases the FSHR intracellular trafficking. **(A)** Experimental design of by-stander BRET between FSHR-Luc8 and a Ypet-fused FYVE motif. **(B)** Time course of BRET assay in HEK293A cells transiently expressing the FSHR-RLuc8, FYVE-Ypet, and either NR or iPRC2. Data are presented as mean ± SEM (N=3). **(C)** Representative image of an FSHR recycling event (indicated by the blue arrow) obtained by TIR-FM in HEK293 cells stably expressing SEP-FSHR and transiently expressing either NR-mCherry or iPRC2-mCherry. **(D)** Number of recycling events of SEP-FSHR per minute and per µm^2^, measured in real time using TIR-FM in HEK293 cells stably expressing SEP-FSHR and transiently expressing either NR-mCherry or iPRC2-mCherry, stimulated with 10 nM FSH. Data represent mean ± SD (n = 11 cells). For each condition, 4 cells were analysed 10 min upon FSH stimulation, and 7 cells 20 min upon 10 nM FSH stimulation, across 3 independent experiments. Statistical significance was assessed by unpaired Mann-Whitney test, ***p < 0.001.

The APPL1 protein plays a role in FSHR recycling and signalling at the VEE, the main recycling compartment of the FSHR. Since the ability of APPL1 to promote the FSHR recycling depends on its phosphorylation status on S410, we sought to investigate whether iPRC2 affects the phosphorylation of APPL1 in response to FSH stimulation. By co-immunoprecipitation and Western blotting (**Figure 6E**), we were able to detect phosphorylation of APPL1 in response to FSH, and the presence of iPRC2 tended to impair APPL1 phosphorylation (**Figure 6F**), although this trend did not reach statistical significance.

## Discussion

In this study, we report the isolation and characterization of an intra-VHH, iPRC2, that alters the FSHR signalling. Importantly, whereas several intra-VHH have served to reveal the sub-cellular sites of activation of several GPCR, iPRC2 induces *per se* a location bias at the FSHR by provoking its accumulation in early endosomes, consequently decreasing its recycling (**Figure 7**).

**Figure 7:**
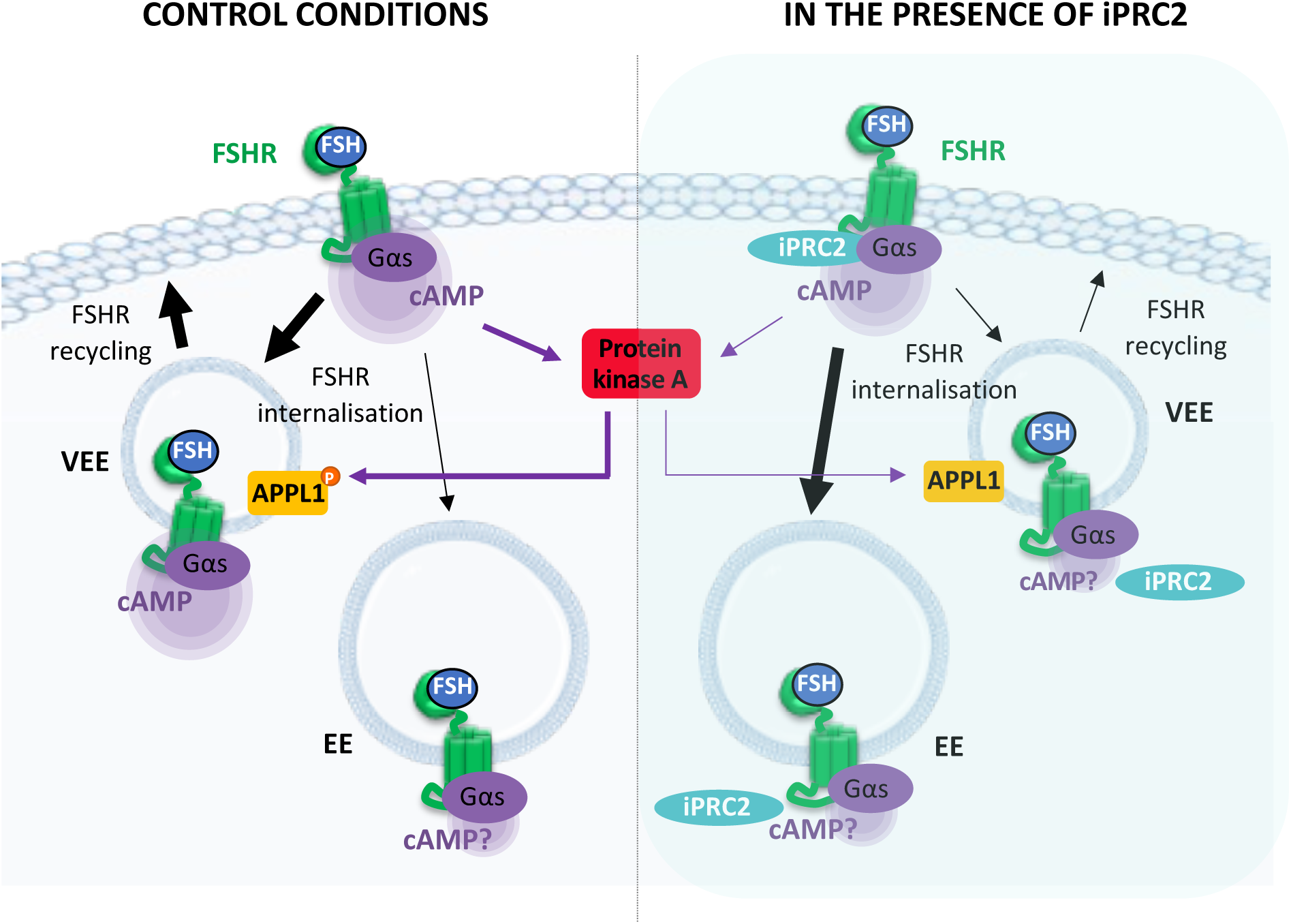
Hypothetical model raised from our data and opened questions. cAMP produced from intra-cellular compartments might be affected by iPRC2 action. By an unknown mechanism, the FSHR accumulates in the EE in the presence of iPRC2, hence less receptor is available to reach the VEE, a prominent recycling component of gonadotropin receptors. This effect could be further enhanced by a diminished PKA activity that would impact APPL1 phosphorylation. It has not been demonstrated in this study whether the FSHR is still associated to G*α*s in the EE.

I*n vitro* experiments indicate that iPRC2 binds not only ICL1 but also ICL3. Since the BBXXB motif in FSHR ICL3 is involved in G protein recruitment to the receptor (51), it was possible that iPRC2 could hinder G*α*s protein binding sterically, which would have explained the inhibition of cAMP and CRE-dependent transcriptional activity by iPRC2. Both readouts depend on G*α*s activation, since G*α*s is responsible for cAMP production, and cAMP activates PKA, which in turn phosphorylates CREB (52,53). Surprisingly, the ectopic expression of Gαs not only failed to displace iPRC2 binding to the FSHR in HEK293 cells depleted for Gαs-q-11/12, but it actually favoured it. This observation, further supported by increased iPRC2 binding to the receptor in parental HEK293 when compared to HEK293 cells depleted for Gαs-q-11/12, suggests that the presence of Gαs enhances iPRC2 binding. This could be due to Gαs inducing a conformation of FSHR more favourable to iPRC2 binding. An intermediate conformation could be stabilized by the iPRC2, as demonstrated in previous studies involving ADRB2 (57). The presence of Gαs would further stabilize a conformation favourable for intra-VHH orientation, which is not visible by binding assay. Another possibility is that iPRC2 stabilizes an interface between ICL1 and 3 and Gαs, thereby increasing the avidity of the intra-VHH when both proteins are present. However, we cannot exclude the possibility that Gαs induces a conformational change that simply improves the energy transfer between the RLuc8 and Venus biosensors, respectively fused the FSHR and to iPRC2. In either case, the fact that iPRC2 recognizes FSHR at comparable levels, whether FSH-activated or not, suggests that Gαs may be pre-coupled to the FSHR prior to stimulation, and could explain why a difference in mini-Gs recruitment to the FSHR is not visible, although one has to keep in mind that mini-G biosensors remain notoriously bound to the receptor once bound (46). Both scenarios align with the selection process on active FSHR, potentially involving Gαs protein.

Gαs activation involves the exchange of GDP for GTP. Even though iPRC2 impairs Gαs -dependent signalling without hindering Gαs binding, no difference in Gαs activation was observed. Interestingly, our experimental setup involves the anchoring of the Gαs-GTP detector to the plasma membrane (47), which selectively measures Gαs activation at this specific cellular location. It is thus likely that iPRC2 affects Gαs -dependent signalling at a different subcellular site, since our results demonstrate that iPRC2 expression alters FSHR intracellular trafficking. Importantly, iPRC2 induced FSHR accumulation in the EE, whereas it had no effect on FSHR internalization, nor on β-arrestin 2 recruitment. As expected from this perturbed trafficking, the receptor recycling was strikingly impaired. The internalized FSHR is recycled through a recently described endosomal compartment, the very early endosomes (VEE) (32). These VEE are smaller than the conventional EE, located closer to the plasma membrane, and devoid of EE and intermediate EE markers such as EE antigen 1 (EEA1), Rab5, and phosphatidylinositol-3 phosphate (PI3P) (31). A subpopulation of the VEE is positive for APPL1. When phosphorylated on its S410 by PKA, APPL1 favours FSHR recycling to the plasma membrane (58). Therefore, we hypothesized that the decrease in cAMP production in the presence of PRC2 could be responsible for lower APPL1 phosphorylation levels, thus diminished recycling. Our data tend to support this possibility, as APPL1 phosphorylation appeared to be reduced in the presence of iPRC2, although the data did not reach statistical significance. Nonetheless, since both APPL1 and iPRC2 interact with FSHR ICL1 (36), recycling could also be impaired by steric hindrance, thus diminishing FSHR availability at the plasma membrane, resulting in lower cAMP production. However, we ruled out this possibility since this is not what we observed in our flow cytometry experiments.

The FSHR signals through Gαs at the VEE, yet it remains unclear whether it also signals at the EE. Increased accumulation of FSHR in the EE may lead to a reduction of FSHR at the VEE, potentially responsible for impaired G protein-dependent signalling. An active Gαs protein detector anchored at the VEE membrane could help clarify this, but currently, no reliable VEE markers are available. The modest reduction in cAMP production in response to FSH observed in the presence of iPRC2 may be due to the portion of FSHR that is redirected to the EE, while signalling from the plasma membrane and VEE remains active. Regardless, we do not yet understand the mechanisms by which iPRC2 redirects a portion of internalized FSHR towards the EE rather than the VEE, and further studies are required to elucidate this process.

It is also important to consider whether iPRC2 promotes FSHR degradation through the lysosomal or proteasomal pathways, which could explain both the decreased recycling and cAMP production, or if iPRC2 inhibits FSHR degradation, leading to its accumulation in the EE. However, again, this would result in a decrease in the total FSHR expression in the cell, which is not consistent with our results.

To our knowledge, iPRC2 is the first identified anti-GPCR intra-VHH inducing *per se* a location bias. Location biases constitute a new level of potential pharmacological regulation. By forcing receptor addressing in selective cell compartments, intra-VHHs like iPRC2 might help deciphering the complexity of GPCR signalling from the plasma membrane or from endosomal compartments, and lead to the development of a new generation of pharmacological tools. Nowadays, the pharmacological consequences of the activation of a GPCR at one place or another are starting to emerge (59,7,8), but the consequences on gene expression have been poorly described. Therefore, iPRC2 is a precious tool to identify the differences in gene expression when the FSHR is retained in the EE and less recycled, especially in a physiologically relevant cell system as primary Sertoli cells of the testis or granulosa cells of the ovary.

## Acknowledgements

The authors would like to acknowledge the Merck company for kindly providing purified human FSH, Pierre Martineau (Institut de Recherche en Cancérologie de Montpellier, France) for kindly providing the pCANTAB 6 phagemid vector and the KM13 phage and for valuable advices, Dr. M.G. Scott (Cochin Institute, Paris, France) for kindly providing the Ypet-β-arrestin 2 plasmid, Mikel Garcia-Marcos for the kind gift of the ONE-GO biosensors, and Silvia Sposini (University of Bordeaux, France) for her kind help in TIRF image acquisition and processing. We are also indebted to the FILM Facility at Imperial College, London, UK.

## Abbreviations

ADRB2: β2 adrenoceptor
APPL1: Adaptor protein containing PH domain
PTB: domain and leucine zipper motif
AVP: Arginine-vasopressin
AVPR2: Arginine-vasopressin 2 receptor
BRET: Bioluminescence resonance energy transfer
BSA: Bovine serum albumin
CDR: Complementarity-Determining Region
CRE: cAMP-responsive element
CXCR3: Chemokine receptor 3
DMEM: Dulbecco’s Modified Eagle Medium
EE: Early endosome
FSH: Follicle-stimulating hormone
FSHR: FSH receptor
Fsk: Forskolin
GPCR: G protein-coupled receptor
GFP: Green fluorescent protein
GRK: G protein-coupled receptor kinase
HEK293: Human Embryonic Kidney 293
HTRF: Homogeneous time-resolved fluorescence
ICL: intracellular loop
ISO: Isoproterenol hydrochloride
LC: Liquid Chromatography
LHCGR: Luteinizing hormone/choriogonadotropin receptor
mCherry: monomeric red fluorescent protein derived from DsRed of Discosoma sea anemones
Mini-Gs protein: minimal engineered GTPase domain of the Gα subunit
mLTC-1: Murine Leydig Tumoral cells
NGS: Next generation sequencing
PKA: Protein kinase A
PTH: Parathyroid hormone
PTHR: Parathyroid hormone receptor
RLuc8: luciferase from the sea pansy Renilla reniformis
SEP: Superecliptic pHluorin
TIR-FM: Total Internal Resonance-Fluorescence Microscopy
TSHR: Thyroid-stimulating hormone
VEE: Very early endosome
Ypet: yellow fluorescent protein derived from Aequorea victoria

