## Supplementary Material for "A single domain intrabody as a novel tool to bias the subcellular trafficking of the follicle-stimulating hormone receptor"

- Supplementary Figures S1 to S6

- Supplementary Table S1

- Supplementary Methods

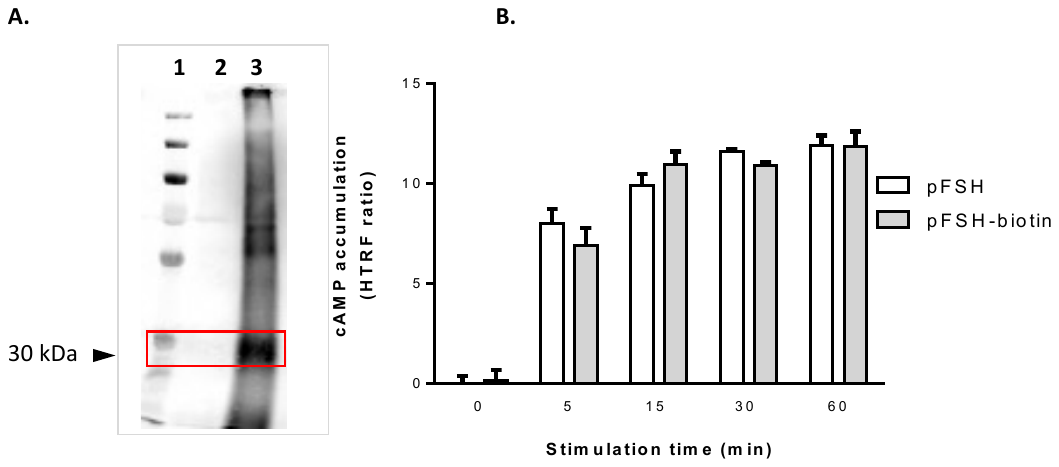

**Figure S1, related to Figure 1. *Control of pFSH biotinylation A.*** Western blot analysis using streptavidine for biotin staining. First lane: molecular weight ladder, second lane: pFSH, third lane: biotinylated pFSH. pFSH expected size is indicated by the red frame. **B.** Comparison of pFSH and pFSH-biotin ability to induce cAMP production, measured by HTRF in HEK293N cells transiently expressing hFSHR after 5, 15, 30 or 60 min of stimulation.

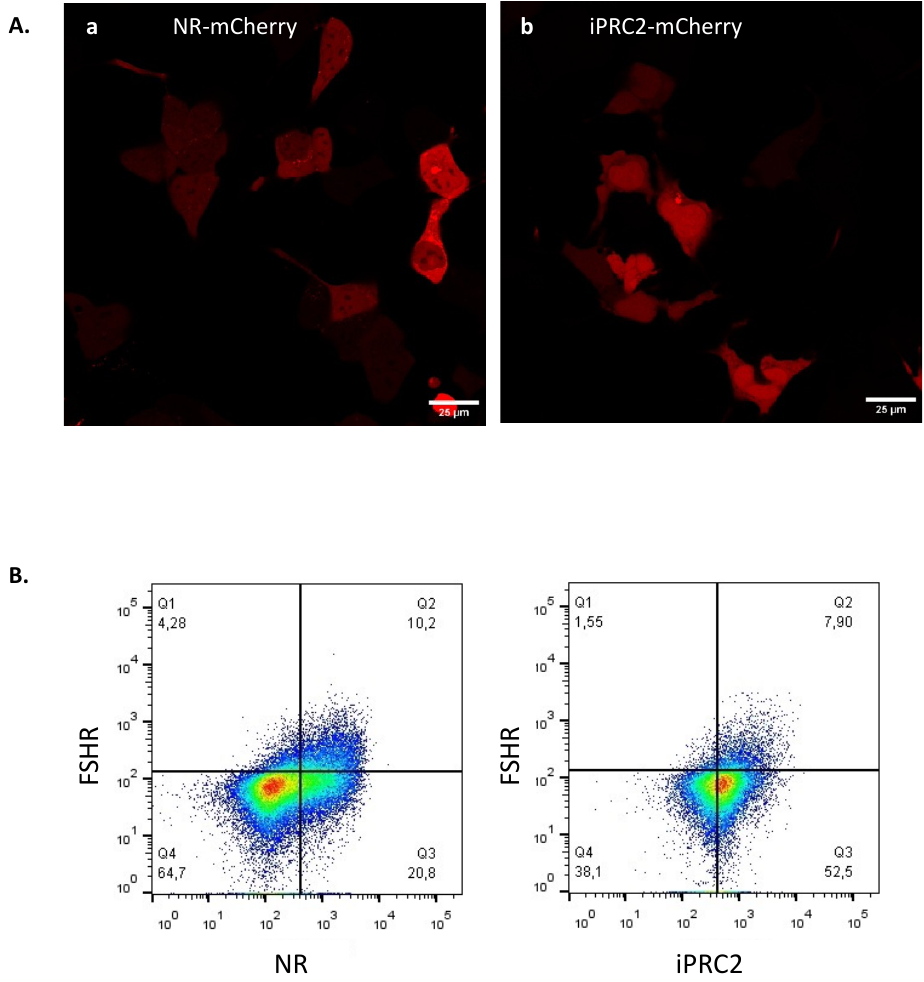

**Figure S2, related to Figures 2, 4 and 6*: iPRC2 and non-relevant (NR) intra-VHH expression in HEK293A cells***. **(A)** Confocal microscopy representative images of NR (a) and iPRC2-mCherry (b) transiently expressed in HEK293A cells. **(B)** Flow cytometry showing the population of cells expressing FLAG-FSHR detected with an anti-FLAG antibody labelled with PE (on the top) and either the irrelevant intra-VHH fused to 6xHis (on the left) or iPRC2-6xHis (on the right), detected with an anti-His_6_ antibody labelled with APC.

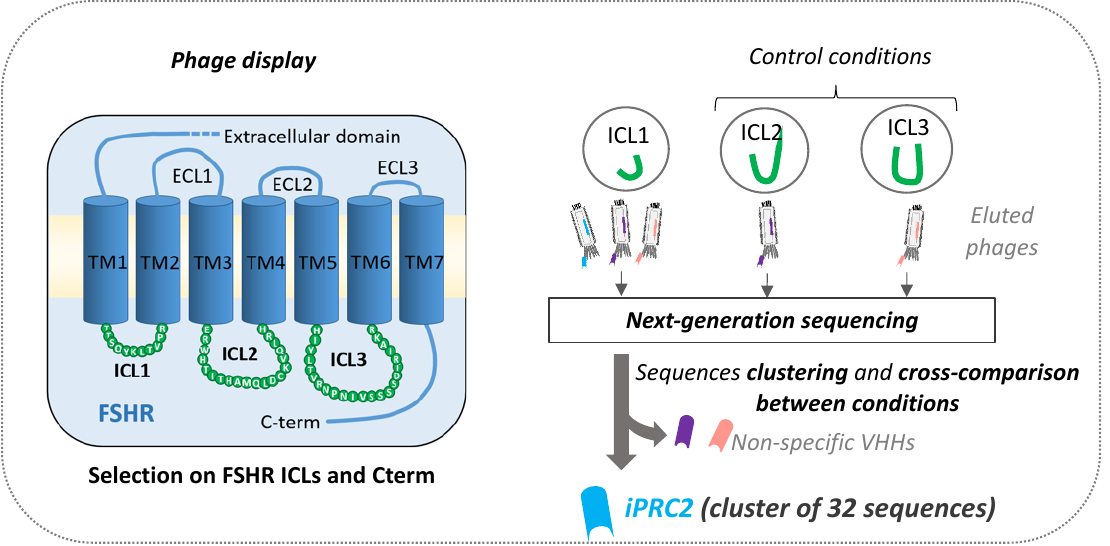

**Figure S3, related to Figure 2G: *VHH selection on FSHR peptides.*** VHH selection was performed on each of the peptides corresponding to the FSHR intracellular loops (ICLs) ICL1, ICL2, ICL3 and on the carboxyterminal region. DNA encoding VHHs from selected phages were sequenced by NGS. The iPRC2 sequence was identified as a specific cluster of 32 sequences/100, 000 sequences selected on ICL1.

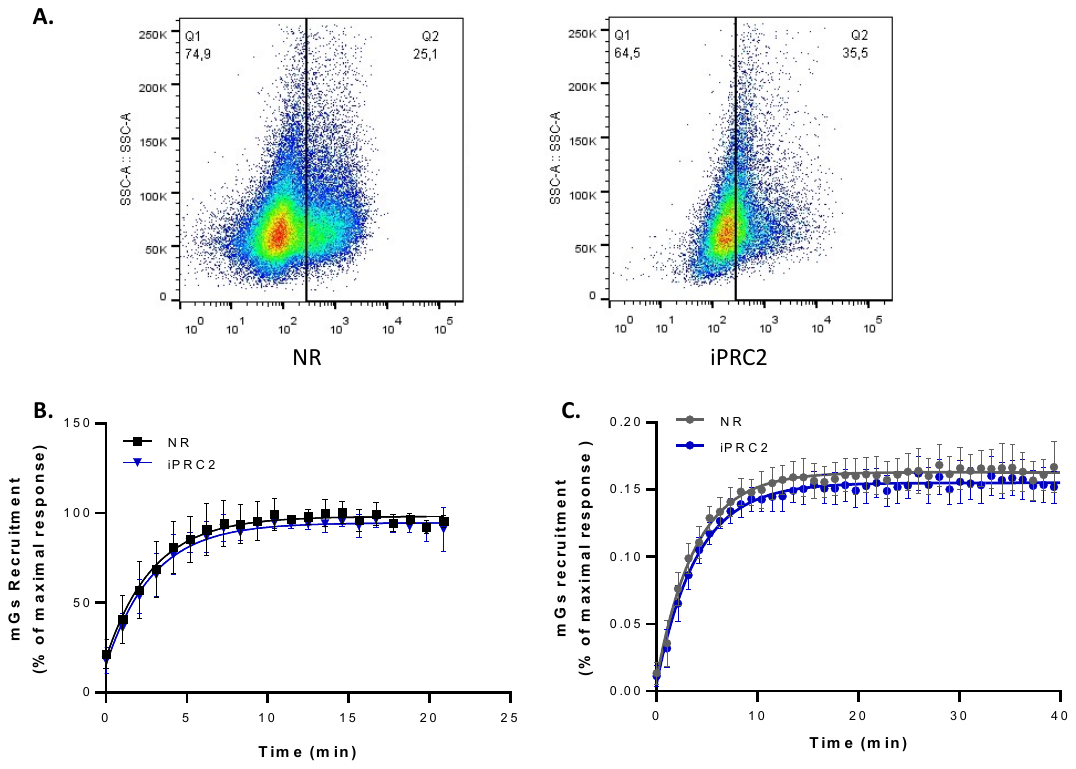

**Figure S4, related to Figure 4*: iPRC2 impairs FSH-stimulated Gs-dependent signaling***. **A.** Flow cytometry showing the population of cells expressing either the irrelevant intra-VHH fused to 6xHis (on the left) or iPRC2-6xHis (on the right). **B.** BRET assay measuring the recruitment of NES-Venus-mGs to FSHR-RLuc8, in the presence of either NR or iPRC2, in response to stimulation with 33 nM FSH of parental HEK293 cells. BRET ratios were expressed as percentage of the maximal FSH-response in the absence of VHH. **C.** Same experiment as (B.) in HEK293 cells depleted of Gαs/q/11/12. Data are represented as mean ± SD, and curves correspond to fitted data to one-phase association curves (N = 4).

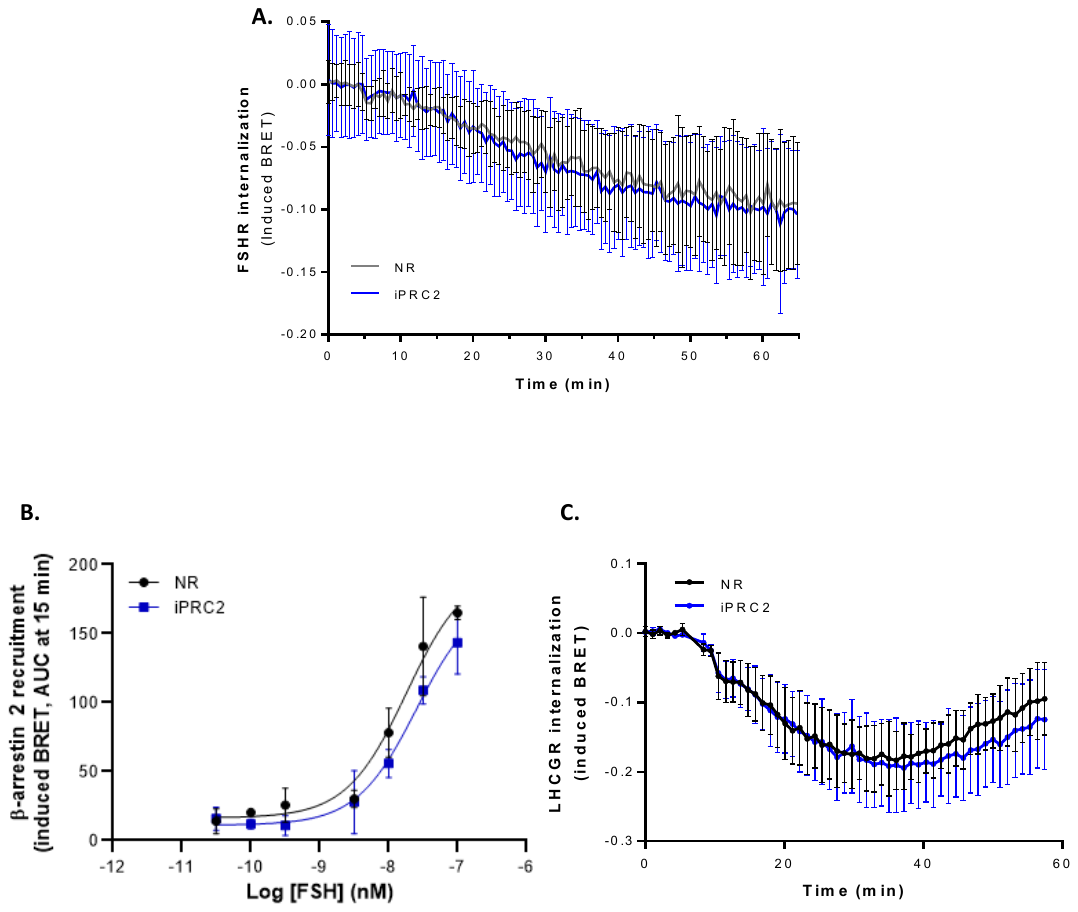

**Figure S5, related to Figure 5.** ***The FSHR internalization remains unaltered in the presence of iPRC2***. **A.** Internalization assay in HEK293A cells transiently expressing the FSHR-RLuc8, and the N-terminus of the fatty acylation motif of the Lyn kinase fused to Ypet and either NR, iPRC2, or empty vector (pCDNA3.1), after stimulation with 30 nM FSH (N=3). **B.** β-arrestin 2-Ypet recruitment to the FSHR-RLuc8 in parental HEK293 cells, in the presence of iPRC2-6xHis or NR fused to 6xHis (N=3). **C.** LHCGR internalization measured by BRET as above, in the presence of iPRC2-6xHis or NR fused to 6xHis (N=3).

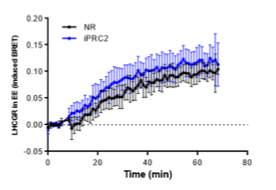

**Figure S6, related to Figure 6.** ***iPRC2 expression does not affect the LHCGR addressing to EE.*** Time course of BRET assay in HEK293A cells transiently expressing the LHCGR-RLuc8, FYVE-Ypet, and either NR or iPRC2. Data are presented as mean ± SEM (N =3).

***Supplementary Material & Methods***

- *Flow Cytometry*

To monitor transient expression, cells were transfected in the same condition as for cAMP measurement: 0.5 ng Flag-tagged human FSHR (43) or 0.25 ng Flag-tagged human AVPR2 (44) plasmid DNA/cm^2^ and 80 ng of VHH-6xHis plasmid DNA/cm^2^ were used for transfection. For the condition without transfected VHH, an empty pcDNA3.1 plasmid was used. Cells were detached following 1 hour-incubation in Hanks Balanced Salt Solution (HBSS) w/o Ca^2+^ and Mg^2+^ (Eurobio, Les Ulis, France), and re-suspended in working buffer (PBS w/o Ca^2+^ and Mg^2+^, 0.5% BSA) 48 h after transfection. Cells were incubated with FcR Blocking Reagent human (1:20, Miltenyi Biotech, Bergisch Gladbach, Germany) for 10 min at 4°C, and with Phycoerythrin-conjugated anti-FLAG antibody (DYKDDDDK Antibody PE, 1:1000, Miltenyi Biotech, Bergisch Gladbach, Germany) and LIVE/DEAD™ Fixable Violet Dead Cell Stain (1:1000, Invitrogen, [Waltham, MA, USA](https://www.google.com/search?client=firefox-b-d&sxsrf=APwXEdewn3ShburY1PA7yl5NGOUDQy5Kvg:1681983220401&q=Waltham&si=AMnBZoFk_ppfOKgdccwTD_PVhdkg37dbl-p8zEtOPijkCaIHMjrOwoPM9hDMB6S9ndin1hg2iYOJftIs7F6s0tPrPWnJTgovlvP3udfl4XdpoUGNfP7s4P_Qj1B2zQFVRpZqKvzU5cNNjldpmAupMVuSmqmih4hA1dsAnmUwlykyQZBBWogIiH-sT_HrYhrkeBR0TUuRn5S4&sa=X&ved=2ahUKEwjF0_rik7j-AhVXQaQEHUvyAU4QmxMoAXoECCMQAw)) for 30 min at 4°C. Cells were washed twice in working buffer before permeabilization using BD Cytofix/Cytoperm™ Fixation/Permeabilization kit (BD Biosciences, Franklin Lakes, NJ, USA) following manufacturer’s protocol. Allophycocyanin-conjugated anti-6xHis antibody (His antibody APC, 1:1000, Miltenyi Biotech, Bergisch Gladbach, Germany) was added for 30 min at 4 °C. Following two washes in perm/wash buffer, cells were resuspended in PBS for analysis with the MACSQuant Analyzer 10 Flow cytometer (Miltenyi Biotech, Bergisch Gladbach, Germany). Data were analyzed and plotted with the FlowJo software (FlowJo, Ashland, OR, USA).

- *Confocal microscopy*

HEK293A cells were seeded on glass chambers coated with Poly-L-Lysine (Sigma-Aldrich, St. Louis, MO, USA), and transfected with 0.3 µg plasmid DNA/cm^2^ of intra-VHH-mCherry. An irrelevant intra-VHH not binding the FSHR (NR), previously described in (41), has been used as a negative control in all following experiments. Forty-eight hours after transfection, cells were washed with DMEM w/o phenol red and images were acquired on a ZEISS Axio Observer Z1 using a 40X oil immersion objective and analysed with the ZEN software (Zeiss, Oberkochen, Germany)***.***
